# Time course of changes in the long latency feedback response parallels the fast process of short term motor adaptation

**DOI:** 10.1101/2020.04.25.061382

**Authors:** Susan K. Coltman, Paul L. Gribble

**Author notes:** Please Address Correspondence to: Paul L. Gribble, Brain and Mind Institute, Dept. of Psychology, Western University, 1151 Richmond St., London, ON, Canada, N6A 3K7.

## Abstract

Adapting to novel dynamics involves modifying both feedforward and feedback control. We investigated whether the motor system alters feedback responses during adaptation to a novel force field in a manner similar to adjustments in feedforward control. We simultaneously tracked the time course of both feedforward and feedback systems via independent probes during a force field adaptation task. Participants (n=35) grasped the handle of a robotic manipulandum and performed reaches to a visual target while the hand and arm were occluded. We introduced an abrupt counter-clockwise velocity-dependent force field during a block of reaching trials. We measured movement kinematics and shoulder and elbow muscle activity with surface EMG electrodes. We tracked the feedback stretch response throughout the task. Using force channel trials we measured overall learning, which was later decomposed into a fast and slow process. We found that the long-latency feedback response (LLFR) was upregulated in the early stages of learning and was correlated with the fast component of feedforward adaptation. The change in feedback response was specific to the long-latency epoch (50-100 ms after muscle stretch) and was observed only in the triceps muscle, which was the muscle required to counter the force field during adaptation. The similarity in time course for the LLFR and the estimated time course of the fast process suggests both are supported by common neural circuits. While some propose that the fast process reflects an explicit strategy, we argue instead that it may be a proxy for the feedback controller.

**New & Noteworthy:** We investigated whether changes in the feedback stretch response were related to the proposed fast and slow processes of motor adaptation. We found that the long latency component of the feedback stretch response was upregulated in the early stages of learning, and the time course was correlated with the fast process. While some propose that the fast process reflects an explicit strategy, we argue instead that it may be a proxy for the feedback controller.

## Introduction

It has been proposed that the brain uses internal models to predict the sensory consequences of a motor command (Desmurget and Grafton, 2000; Miall and Wolpert, 1996; Wolpert et al., 1998; Wolpert and Flanagan, 2001). The use of predicted sensory consequences reduces movement instabilities arising from delays in the actual sensory feedback (Miall and Wolpert, 1996; Wolpert et al., 1998; Wolpert and Kawato, 1998). The driving force of motor adaptation is sensory prediction error, which results from the mismatch between the predicted consequences of a motor command and sensory feedback.

The ability to adapt our movements in response to changes in both our body and the environment is critical for maintaining accurate motor performance. When a movement being performed results in an unexpected outcome, this generates an error signal. Error-based models of motor learning have been shown to fit trial to trial adaptation to perturbations extremely well (Smith et al., 2006; Thoroughman and Shadmehr, 2000). The use of state-space models suggests that a portion of the error experienced can be corrected for trial to trial, referred to as the rate of learning, while the passage of time, in the absence of error, leads to forgetting (Smith et al., 2006; Thoroughman and Shadmehr, 2000). Several recent studies have proposed various novel methods to examine the simultaneous operation and interaction of multiple learning processes (Cheng and Sabes, 2006; Kording et al., 2007; Smith et al., 2006).

A prominent model for error-based learning, proposed by Smith et al. (2006), suggests that the experience of error engages at least two independent processes, a fast and a slow process, where overall learning during short term adaptation will be the sum of these two processes. Mathematically, both the fast and slow processes have the same form, except that the learning rate and retention parameters are different for each process: the fast process learns much more quickly than the slow process, however the fast process forgets more rapidly than the slow process.

Recent evidence provided an alternative decomposition of overall learning during visuomotor rotation tasks into an explicit aiming strategy and an implicit process (Taylor et al., 2014). It has since been suggested that the rapid initial improvement is due to learning in the fast process that compares to the explicit process, whereas the gradual learning that follows is due to a slow process that compares to the implicit process (McDougle et al., 2015). However, the use of an explicit strategy during force field adaptation remains open to debate.

At an early stage of learning novel dynamics, such as force field learning, changes in muscle activity are mainly due to corrective feedback responses (Thoroughman and Shadmehr, 1999). When observing the visuomotor feedback gains over the course of an adaptation task, Franklin et al. (2012) demonstrated that both the introduction and the removal (after adaptation is complete) of a velocity-dependent force field modified the magnitude of the gain of the feedback response. This finding supports the understanding that feedback responses are used to restore stability against unexpected movement outcomes. These feedback contributions to the overall motor command are then reduced as feedforward control is learned.

Franklin et al. (2012) suggested that there was a relationship between error size and the visuomotor feedback response. Previous research has shown that learning from larger errors engages a different process than learning from smaller errors, based on experimental data comparing abrupt versus gradual perturbation schedules (Orban de Xivry et al., 2011; Schlerf et al., 2012; Tseng et al., 2007). Within the framework of a two-state model of adaptation, the output of the fast process contributes significantly more than the slow process to overall output during early learning of abrupt perturbations, when errors are the largest. Moreover, the gradual shift from a feedback-driven mode of control to more predictive, feedforward control, reported by Thoroughman and Shadmehr (2000) in response to a novel force field perturbation could be seen as the interaction of the fast and slow processes over the course of learning. Crevecoeur and colleagues (Crevecoeur et al., 2020a, 2020b) recently provided support for adapting to error within a single reaching trial, which leads to the question of whether reflexes are also adapting.

While Franklin et al. (2012) focused on the visuomotor response, error corrections during movement also arise from muscle stretch responses. In recent work Scott (2012, 2004) proposes that the majority of the sophistication embodied in the feedback stretch response occurs during the long latency component and not the short latency component. Recently, Ahmadi-Pajouh and colleagues (2012) measured the long-latency feedback response (LLFR) before and after adaptation to a velocity-dependent curl field and reported an increase in LLFR post-adaptation. Subsequently, Cluff and Scott (2013) measured the LLFR as participants adapted their reaching movements to a velocity-dependent force that resisted elbow motion. Similar to Ahmadi-Pajouh et al. (2012), they showed an increase in LLFR over the course of learning. The results of these two studies suggested that the time course of the LLFR may relate to the slow process in a two-state model of motor adaptation. This is in contrast to the results described above that suggest that the time course of learning-related changes in the visuomotor feedback response relates to the estimated fast process.

In the present study we tracked the time course of the feedback stretch response via randomly interspersed force probe trials during a force field adaptation task in which participants produced point to point reaching movements with the upper limb. The goal was to characterize the time course of changes in the LLFR and directly compare it to the time course of the fast and slow processes of learning, estimated using a two-state model (Smith et al., 2006; Coltman et al., 2019). We found that the time course of the LLFR was highly correlated with the time course of the estimated fast process. Our study sheds light on the debate about the origins of the fast and slow processes during adaptation. Our results demonstrate that the fast process parallels the modulation in gain of the feedback response over the course of learning. We are interested in the question of how the internal estimate of the dynamics of the environment is formed and used for control. We propose that the fast process, estimated from overall learning during force field tasks, may alternatively be an identification of the feedback controller, while the slow process is the recalibrated forward model. Additionally, we discuss the ways in which the LLFR and the fast component may be organized to support motor adaptation by learning from errors.

## Materials and Methods

### Participants

A total of 55 healthy young adults (18 - 34 years of age; 36 females) participated in a force field adaptation experiment. All participants were recruited from the research participation pool maintained by the Department of Psychology at Western University and received either course credit or CAD$18.00 for participation. All participants self-reported being right-handed and had normal or corrected-to-normal vision. The protocol was approved by Western University’s Research Ethics Board and all participants signed a written consent form.

### Apparatus and experimental task

Participants were instructed to make point-to-point reaching movements in a horizontal plane using the KINARM endpoint robot (Kingston, Ontario). Visual information was projected in a horizontal plane at the same level of the hand, via a liquid crystal display monitor and a semi-silvered mirror. Direct vision of the upper limb was blocked using a physical barrier. Participants’ right forearm was supported against gravity by a lightweight sled. Air jets in the sled reduced friction between the sled and the tabletop as participants moved their arm.

A circular cursor (6.5 mm radius) was displayed on the semi-silvered mirror and was used to represent the position of the centre of the robot handle. Participants were presented with a circular start position (6.5 mm diameter) and a circular target (10 mm diameter) located 15 cm away from the start position, 45° to the left of the participant’s midline (**Fig. 1A**). When the cursor entered the start position, a 3.5 N background force was applied in a direction that was 90° CCW to a line joining the start position and the target. This corresponded to the direction of the CCW force field presented in the adaptation block. The purpose of the background force was to ensure that the pre-movement state of the triceps muscle remained consistent across all phases of the experiment, so that changes in feedback responses were comparable. The background force was ramped up over 500 ms, and remained on until participants arrived at the target. Critically, the background force was active on all trials so that participants could not predict the occurrence of feedback probe trials.

**Figure 1.**
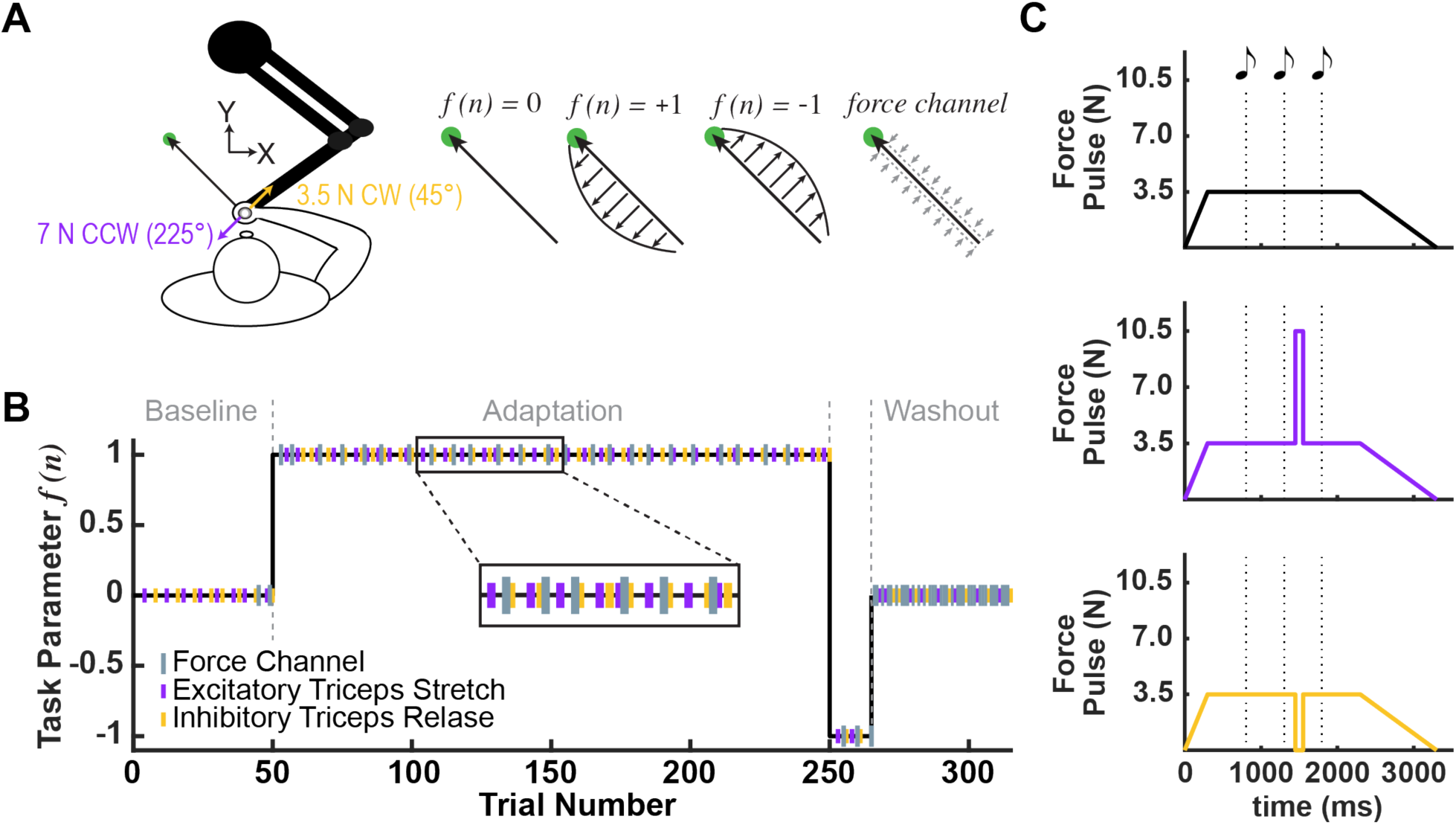
Experiment 1. A, participants held the handle of a robotic manipulandum and performed reaching movements to a visual target (shown in green). Yellow and purple arrows represent the strength and direction of the force pulse used to probe the feedback response. B, Force channel trials (gray bars) were used to track the progression of feedforward control. Force channel trials were randomly distributed during baseline and adaptation blocks and throughout the last block in each session. Force probes (yellow and purple bars) were used to track the change in the feedback response. Force probes were randomly distributed during all phases of testing. C, On each trial, participants heard three tones (100 ms in duration, separated by 500 ms), and were instructed to initiate their reach with the third tone. On each trial the background force was applied, ramped up over 500 ms. The background force remained on until participants successfully arrived at the target. The background force was ramped down over 1000 ms. On random trials, we probed the feedback response using a 100 ms duration, 3.5 N CW force pulse (shown in yellow), or a 7 N CCW (shown in purple) force pulse. Non-probe trials are shown in black.

After maintaining the cursor in the start position for 500 ms, participants were presented with three auditory tones, each 100 ms in duration, and each separated by 500 ms. Participants were instructed to initiate their reach coincident with the third tone (**Fig. 1C**). When the third tone was played, the target changed colour from white to green, representing a secondary “go” signal for participants to initiate movement.

At the end of each movement, participants were provided with feedback indicating the time taken to reach the target. Participants were instructed to bring the center of the handle within 5 mm of the target within 350 - 500 ms. Target colour changed to reflect movement speed: red for movements that were too fast (< 350 ms), blue for movements that were too slow (> 500 ms). The target remained green to indicate that the movement was within the desired timing window. Feedback related to movement time was displayed on the screen for 1000 ms, while the background force was ramped off. The robotic arm then returned the participant’s hand to the start position. Participants were instructed to try to obtain the “good” feedback as often as possible throughout the experiment.

### Experiment 1

We collected data from 35 participants (18 - 34 years of age; 23 females). On a given trial, during the reaching movement the manipulandum either applied no force, a CCW force field, a clockwise (CW) force field, or a force channel. The force field was introduced abruptly on trial 51 (see below) and was defined as:

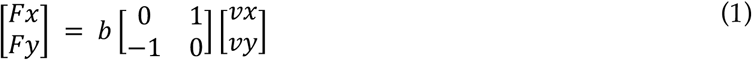

where x and y are lateral and sagittal directions, Fx and Fy are robot forces applied at the hand, vx and vy are hand velocities, and b is the force field constant (± 15 N·s·m^-1^). The sign of the force field constant b determined the direction of the force field (positive = CW and negative = CCW). The effect of the force field for linear movements is to perturb the hand in a direction perpendicular to the instantaneous direction of movement, with a force proportional to movement speed (**Fig. 1A**).

During force channel trials the robot motors were used to constrain movements to a straight path connecting the start position and target (Scheidt et al., 2000; Smith et al., 2006). The channel was approximately 1 mm wide, and the stiffness of the walls was 6,000 N/m with a damping coefficient of 50 N·s·m^-1^. The use of force channels allowed for the removal of kinematic movement errors by effectively preventing any motion perpendicular to the target direction. Without this error signal to drive corrective responses, we assume the output during a force channel is a proxy for the descending motor commands being generated according to the plan desired kinematics and predicted dynamics. Therefore, force channel trials allowed us to probe the feedforward system during learning by measuring the lateral forces exerted on the channel walls.

The experimental session included 315 trials and was divided into four blocks: baseline trials, force field adaptation, brief force field reversal, and finally a series of force channel trials (Coltman et al., 2019; Smith et al., 2006). The first 50 trials were baseline trials during which the robot applied no forces other than the constant background load (see above). On trials 51-250 participants adapted to a CCW force field, followed by a brief force field reversal for trials 251-265. On trials 266-315, a force channel was introduced. In addition the force channel was present for 2 baseline trials (trials 45, 49) and 24 adaptation trials (trials *45, 49, 53, 57, 67, 75, 83, 89, 99, 107, 115, 121, 131, 139, 149, 155, 165, 171, 179, 185, 193, 201, 211, 217, 227*, and *235*).

We probed the feedback response via a 100 ms duration, 3.5 N force pulse aligned with the direction of the CW field (shown in yellow; **Fig. 1A and C**), or a 7 N force pulse aligned with the direction of the CCW field (shown in purple; **Fig. 1A and C**). Feedback probes were applied during 100 of the 315 trials in the task (50 for each force pulse direction). Feedback probes were designed to excite or inhibit the triceps lateral head during the preparatory period before reach onset, at 350 ms before the third tone (Ahmadi-Pajouh et al., 2012). Feedback probes alternated between excitation or inhibition of the triceps and occurred every 2 or 4 trials (see **Fig. 1B**). The force field and force channels were present only during the reach and not in the preparatory period. This meant that the pulses were delivered under identical conditions, regardless of the trial type.

The force pulses displaced the participant’s hand outside the start position. Participants were instructed to try to bring the cursor back to the start position. They were told that whether they were in the start position or not, they should initiate movement to the reach target coincident with the third tone. The order of feedback probe trials, force channel trials, and reaching trials was pseudo randomized so that each participant saw the same order of trials, but a given trial type was not predictable. Additionally, a force channel or force pulse never occurred on the same trial.

### Kinematic Data Analysis

Position, velocity, and force at the handle of the robotic arm were sampled at 1000 Hz. The data were low-pass filtered at 15 Hz. We performed data analysis using custom MATLAB (r2018b, Mathworks) scripts.

For each reaching trial we computed the perpendicular deviation at peak velocity of the hand path relative to a straight line between the start position and target. We then analysed the lateral forces that participants generated throughout movements in the force channel trials. As a measure of the degree of adaptation in force channel trials we computed an adaptation index by estimating the slope of the relationship between the measured lateral force profile produced by the hand (while velocity exceeded 2 cm/s) and the ideal force profile. A linear model with zero intercept was used to estimate the adaptation index on each force channel trial. The ideal force profile was calculated as the force profile that would have to be generated to fully compensate for the force field throughout the movement, had the force field been applied (see Smith et al. 2006 for more details). If these force profiles were uncorrelated, the adaptation index was zero; if these force profiles were identical to one another, the adaptation index was one.

### Model fitting

Smith et al. (2006) described a two-state model of motor adaptation in which overall learning can be decomposed into two separate processes: a fast process xf that learns quickly but has poor retention (Eq. 2) and a slow process xs that has better retention but learns more slowly (Eq. 3). Retention is characterized by the parameters Af and As, for the fast and slow processes respectively. Learning rate is characterized by Bf and Bs. The two processes are combined to produce net motor output xnet (*Eq. 4*). Error e arises on each trial n when there is a difference between the net output xnet and the task parameter f (e.g. the strength of the force field) (*Eq. 5*).

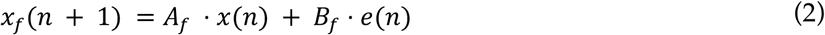

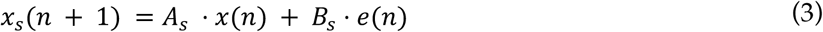

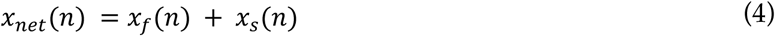

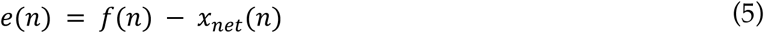

To estimate the retention parameters Af and As and learning rates Bf and Bs we fit the model to the experimental data (using the function *fmincon* in MATLAB r2018b) by minimizing the squared difference between the model predicted net output (*xnet*) and average participant adaptation index, measured on force channel trials. A constrained parameter space was defined by linear inequality constraints and upper/lower bounds (Albert and Shadmehr, 2018). Linear inequality constraints were specified to enforce traditional two-state model dynamics according to:

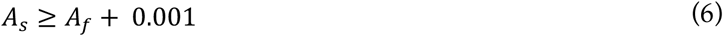

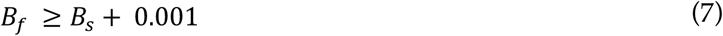

### Electromyographic recordings and analysis

Muscle activity from upper limb muscles was recorded using bipolar surface electrodes (Delsys Bagnoli-8 system with DE-2.1 sensors). EMG signals were amplified (gain = 10^3^) and sampled at 1000 Hz from biceps brachii, lateral head of the triceps, pectoralis major, and posterior deltoid. Prior to electrode placement we cleaned the skin with rubbing alcohol, and electrode contacts were coated with conductive gel. Electrodes were attached to the skin using double-sided adhesive stickers. A ground electrode was placed over the participant’s left clavicle.

On feedback probe trials EMG data were aligned to the initiation of the force pulse, and on reaching trials EMG data were aligned 300 ms prior to movement onset (defined when tangential velocity > 0.5 cm/s) (Cluff and Scott, 2013). The data were then bandpass filtered between 20 Hz and 450 Hz (second-order, dual pass Butterworth filter), full-wave rectified and normalized. Participants were required to hold the cursor within the start position for 500 ms to initiate the three tones, and simultaneously the trial. On each trial, EMG was normalized (per muscle, per participant) to the average EMG activity between 400 ms and 100 ms prior to the first tone. EMG over the whole trial was divided by this value such that a value of 1 represents the mean activity of a given muscle when countering a constant 3.5 N CCW force (Maeda et al., 2018; Pruszynski et al., 2008).

To test for changes in the short and long latency components of the stretch response of the triceps lateral muscle during adaptation, we computed mean EMG over previously defined epochs (Pruszynski et al., 2008); we calculated mean EMG during a pre-pulse epoch (PRE: -50 - 0 ms), short-latency epoch (SLFR: 25 - 50 ms), and long-latency epoch (LLFR: 50 - 100 ms). To obtain a single measure of the LLFR and to track this over the course of learning we subtracted EMG measured during inhibition from EMG measured during excitation. The two types of force pulses (excitation vs inhibition) alternated throughout the task (See **Fig. 1B**), so the difference was taken between each pair of probes. We then averaged this difference signal between the window of 50 - 100 ms to generate a single value, termed the delta LLFR. The values of delta LLFR were then aligned with the feedback probes trials that excited the triceps. The use of delta LLFR versus the LLFR recorded on the trials where we only excite the triceps allows us to infer that the response is directionally independent.

### Control experiment

Here we leverage a technique in which we introduce a force field perturbation gradually over many trials to further test the idea that modulation of the LLFR is dependent upon experiencing large movement errors. Previous research suggests that learning from smaller errors engages a different process than learning from large errors that occur during abrupt perturbations schedules (Criscimagna-Hemminger et al., 2010; Izawa et al., 2012; Schlerf et al., 2012; Tseng et al., 2007). We collected data from 20 participants (18 - 22 years of age; 13 females). All features of the task remained the same, with the exception that the force field was introduced gradually. The strength of the field was increased linearly over 200 trials and then held at a fixed strength, the same strength as the force field introduced abruptly in Experiment 1 (15 N·s·m^-1^), for another 15 trials (**Fig. 9A**). By introducing the perturbation gradually participants reached in the same environment after the perturbation was fully ramped on, while only experiencing small errors from trial to trial during adaptation.

**Figure 2.**
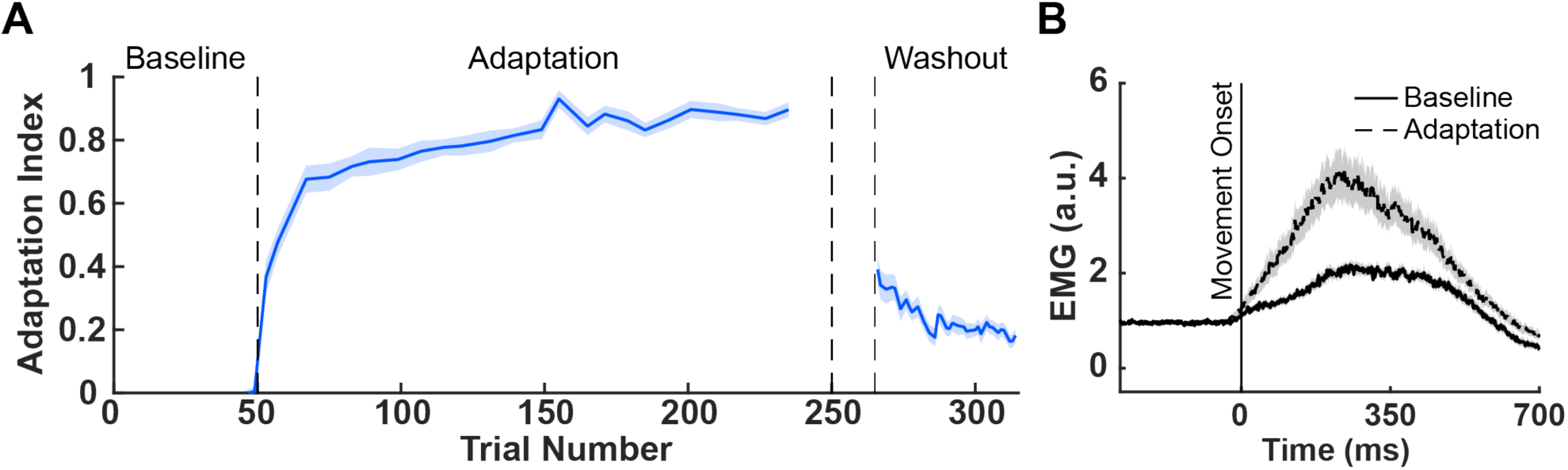
A, The across participant average adaptation index for all force channel trials. The shaded region denotes SEM. B, Triceps EMG during movement increased after adaptation to a CCW force field. EMG time series for non-probe trials averaged over participants (mean ± SEM), is shown during baseline (solid black line) and the second half of the adaptation block (dashed black line).

**Figure 3.**
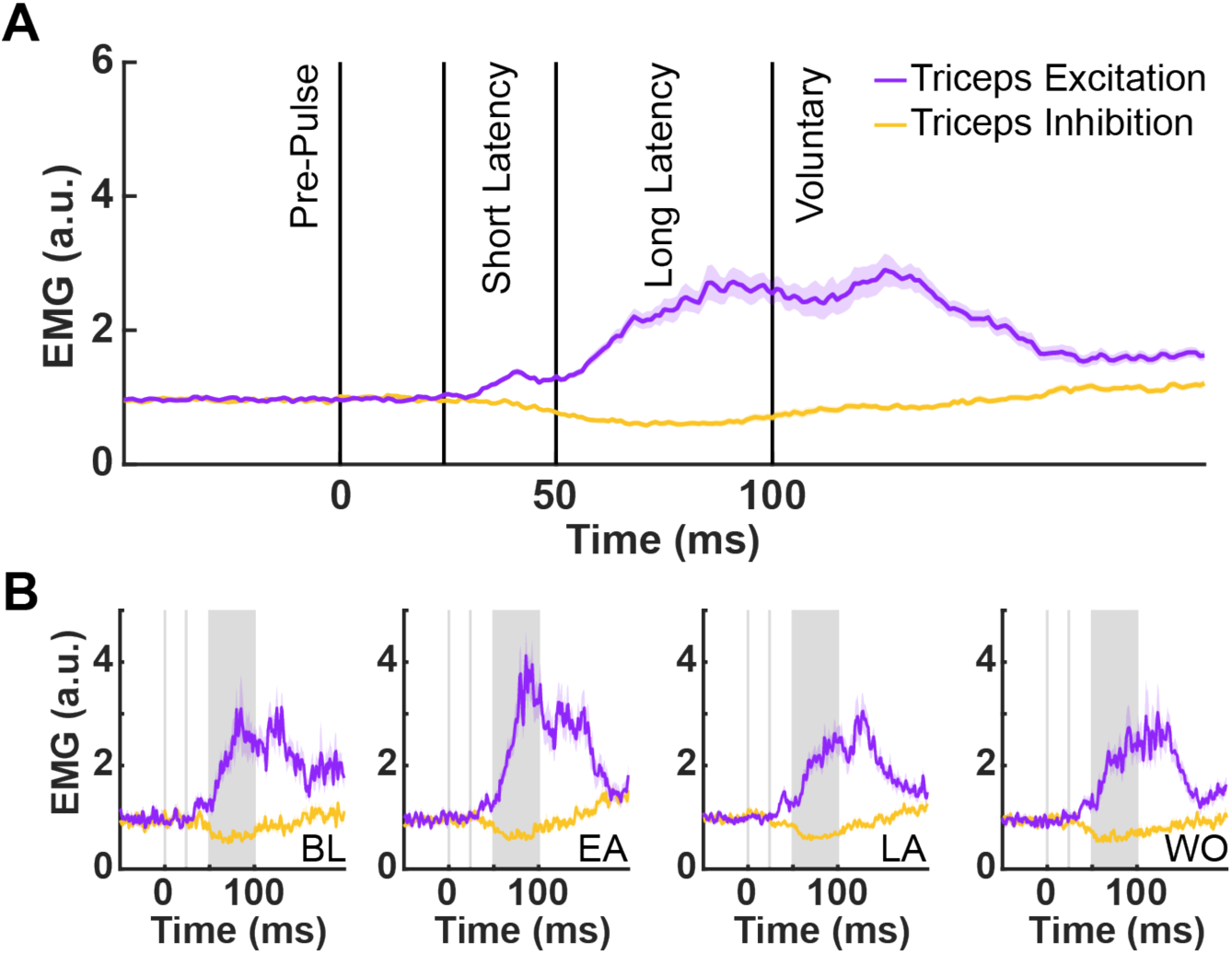
A, Triceps EMG following force pulse probe trials, averaged across all phases of learning and all participants, for trials in which the triceps was stretched (purple) or released (yellow). Data are aligned to force pulse onset (time = 0 ms), and solid vertical lines indicate the different phases of the feedback response. B, Triceps EMG following force pulse probe trials during four phases of learning (BL = baseline, EA = early adaptation, LA = late adaptation, and WO = washout).

**Figure 4.**
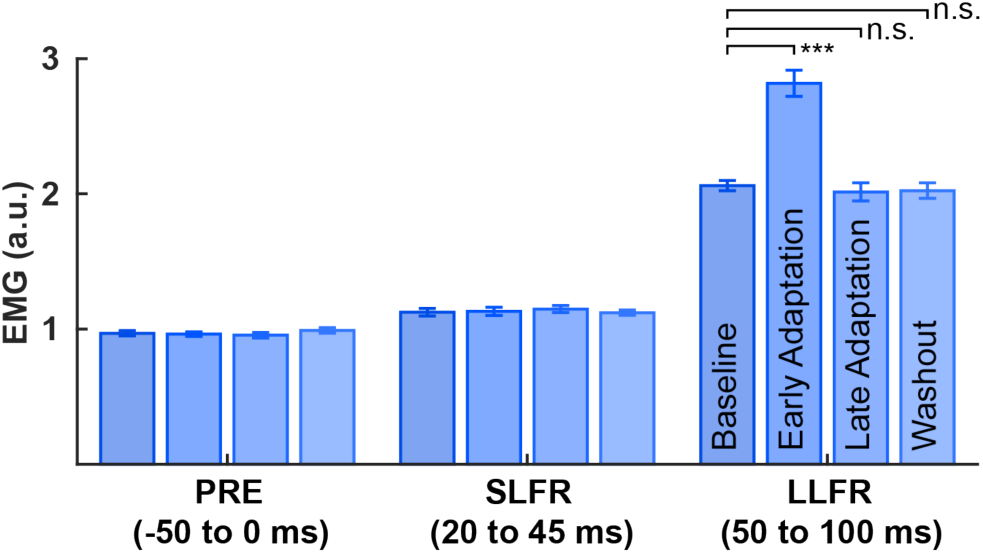
Mean EMG response to force pulse trials across four phases of the adaptation paradigm, and binned in time to reflect the different epochs of the stretch response. Error bars indicate within-subject SEM.

**Figure 5.**
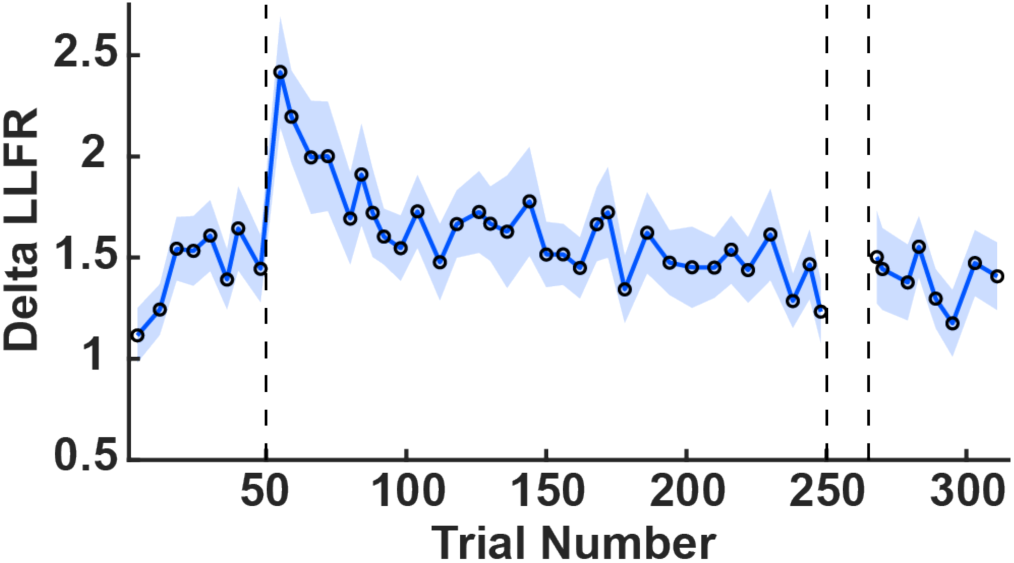
Time course of the triceps LLFR. A, The average change in the LLFR across participants, aligned to trial number for probes trials that stretched the triceps. The shaded region denotes SEM. The ∘ are used to highlight the data points.

**Figure 6.**
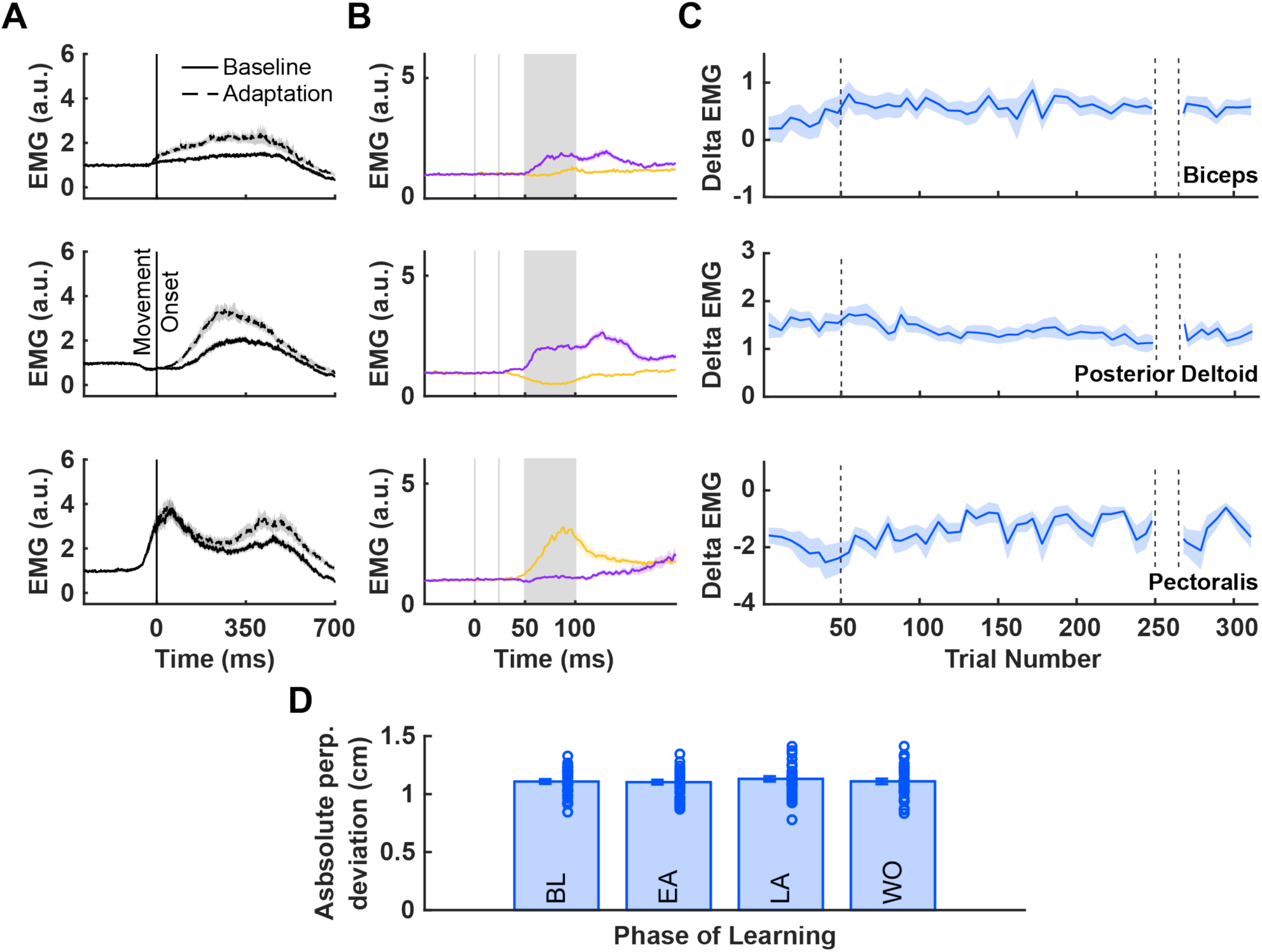
A, EMG time series during arm movement for non-probe trials averaged over participants (mean ± SEM), is shown during baseline (solid black line) and the second half of the adaptation block (dashed black line) for biceps, posterior deltoid and pectoralis muscles. B, EMG during feedback probe trials for biceps, posterior deltoid and pectoralis muscles following force pulse probes, averaged across all phases of learning and all participants, for trials in which the triceps was stretched (purple) or released (yellow). Data are aligned to force pulse onset (time = 0 ms), and solid vertical lines indicate the different phases of the feedback response. C, Time course of changes in EMG for biceps, posterior deltoid and pectoralis muscles, averaged across participants. The shaded region denotes SEM. D, Mean perpendicular displacement of the hand, between 50 to 100 ms after force pulse onset, as a function of learning phase (BL = baseline, EA = early adaptation, LA = late adaptation, and WO = washout). Each circle represents one participant.

**Figure 7.**
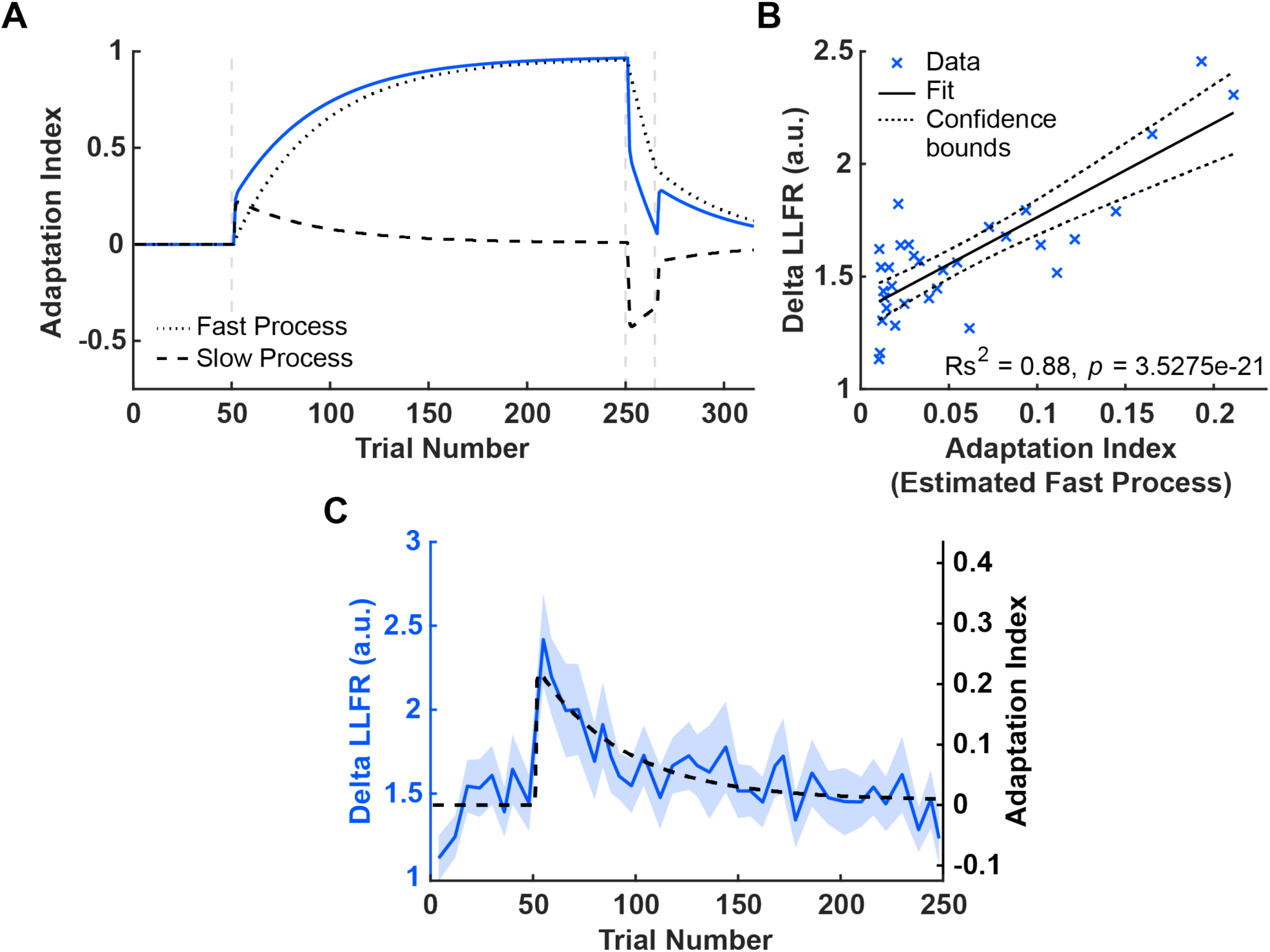
Relating feedforward and feedback control. A, Model simulation. The overall output predicted by the model is shown as a solid blue line. The fast process (dotted black line) and the slow process (dashed black line) are shown as a function of trial. B, Correlation between the mean value, averaged across participants, of the delta LLFR during the adaptation phase, averaged across participants, and the corresponding value of the estimated fast process at each trial at which the delta LLFR is measured during the adaptation phase. C, The time course of the fast process (estimated from the feedforward probes, shown as a black dashed line), overlaid on the time course of the measured delta LLFR (shown as blue solid line). The shaded region denotes SEM.

**Figure 8.**
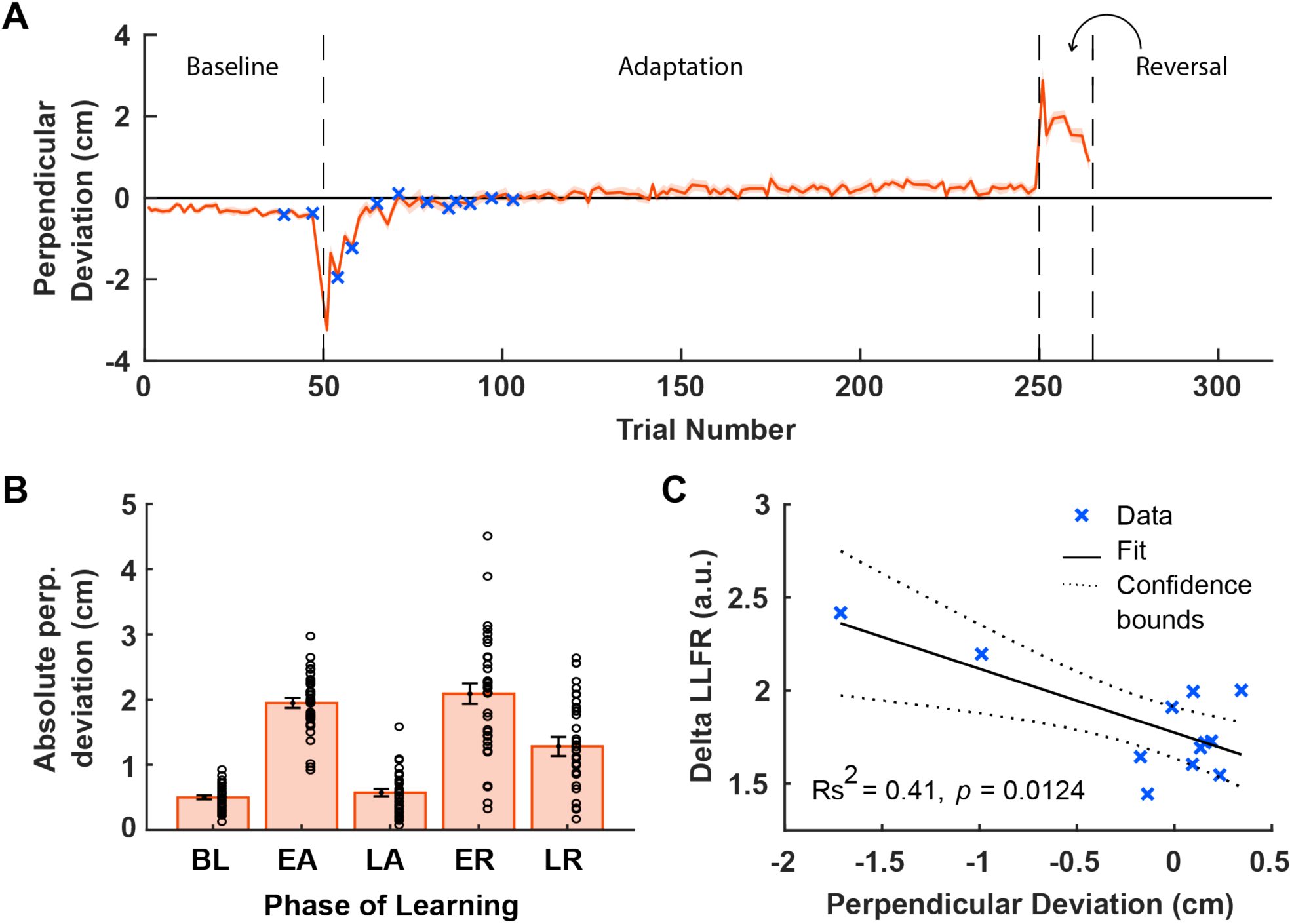
Relating feedback to movement error. A, Average perpendicular deviation of the hand path measured at peak velocity, as a function of adaptation trial. The shaded region denotes SEM. B, Comparison of the mean absolute value of perpendicular deviation for the 4 trials in each phase of learning. Circles represent individual participants. C, Correlation between the delta LLFR and perpendicular deviation of the hand, for 12 trials near the beginning of adaptation (shown in A using a blue x). Dashed lines indicate 95% CI.

**Figure 9.**
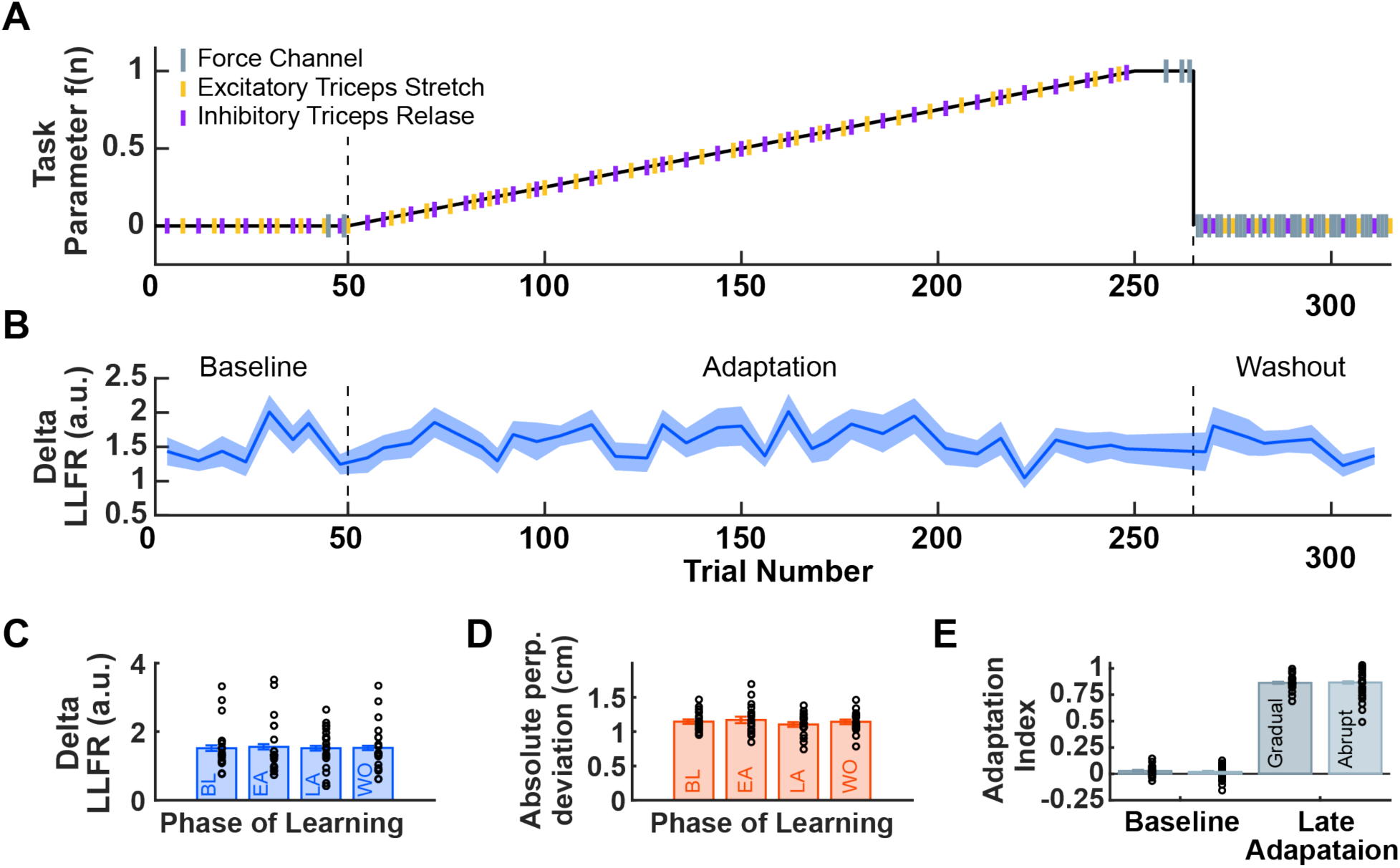
Control Experiment. A, During the adaptation block, the manipulandum applied a CCW force field gradually over 200 trials. This was followed by 15 trials during which the force field was held at a fixed strength. Force probes (yellow and purple bars) were used to track the change in the feedback response in the triceps muscle. B, Average delta LLFR measured in triceps, over the course of learning. The shaded region denotes SEM. C, Comparison of mean perpendicular hand displacement produced by the force pulses as a function of the phase of learning (BL = baseline, EA = early adaptation, LA = late adaptation, and WO = washout). Circles represent individual participants. D, Mean absolute perpendicular deviation for the 4 trials in each phase of learning. Circles represent individual participants. E, Mean adaptation index for gradual vs abrupt introduction of force field, averaged across participants, for the 2 force channel trials in baseline, and the last 3 force channel trials at the end of the adaptation block. Circles represent individual participants.

### Statistical analysis

To test differences between means we used within-subject ANOVAs. We used linear regression to quantify the relationship between the changes in the LLFR and the estimated fast process of feedforward adaptation. Pairwise comparisons were performed with paired t-tests. For statistical analyses that require multiple comparisons, we used the Holm-Bonferroni correction (Holm, 1979). Statistical analyses were performed using MATLAB (r2018b, Mathworks). Statistical tests were considered significant at p < 0.05. For all reported values we report the mean and standard error of the mean (SEM).

## Results

Guided by a two-state model of short-term motor adaptation, we investigated whether the time course of changes in the feedback stretch response would resemble the fast process of the feedforward component of adaptation, or the slow process, or neither. In Experiment 1 we incorporated probe trials with force pulses to measure the time course of the feedback stretch response during short-term learning. We used force-channel trials to probe the time course of the feedforward system.

Participants reached to a target 45° left of their midline. We introduced an abrupt CCW velocity-dependent force field during an adaptation block (200 trials). **Figure 2A** shows adaptation index for all force channel trials, averaged across participants. The adaptation index represents the proportion of compensation for the experienced force field. To assess performance after adaptation to the CCW force field we compared the mean adaptation index averaged across participants in the last four adaptation trials, to the mean adaptation index in the two force channel trials in the baseline period before the introduction of the force field. There was a reliable increase in the mean adaptation index at the end of adaptation (0.9026 ± 0.0133) compared to baseline (0 ± 0.0105; t(34) = 49.0160, p = 1.9713e-33), thus demonstrating that participants reliably adapted to the CCW force field.

The EMG data from reaching trials were aligned to 300 ms prior to movement onset, which was defined as the time at which tangential velocity exceeded 0.5 cm/s. **Figure 2B** shows a summary of triceps EMG activity during baseline trials (32 non-probe trials) and the last half of the adaptation block (56 non-probe trials). The Figure shows the across-participant mean EMG (± SEM) of the triceps lateral head for reaching trials in the absence of force pulses or force channels. Because the data have been normalized to the background load applied in the start position, the values of EMG start at 1. Note that the background load remained on throughout the movement, until participants arrived at the target, and this presumably influenced muscle activity in the triceps even in the absence of the force field. We observed an increase in triceps EMG activity beginning near movement onset, after participants adapted to the CCW force field, compared to EMG activity in baseline trials. This shows that in order to compensate for the force field and move straight to the target, participants increased the activation of triceps. Thus in the analyses below we focus on changes in the feedback stretch response of triceps, as this was the muscle primarily involved in producing compensatory force when participants adapted to the CCW force field.

Feedback probes were designed to inhibit or excite the triceps during the preparatory period before reach onset, and were delivered at 350 ms before the third tone (Ahmadi-Pajouh et al., 2012). **Figure 3A** summarizes the feedback stretch response of triceps resulting from excitation (purple line) or inhibition (yellow line). The Figure shows the across-participant average EMG (± SEM) of the triceps lateral head, averaged across all force pulse trials in the experiment. Because the data have all been normalized to the background load applied in the start position, the values of EMG start at 1. The triceps shows both a classic short latency stretch response (25-50 ms) as well as a long latency feedback response (50-100 ms). **Figure 3B** summarizes the feedback stretch response of triceps during four defined phases of learning: the last four force pulse trials during baseline, the first and last four force pulse trials during adaptation (early adaptation and late adaptation, respectively), and the first four force pulse trials in washout. This qualitatively demonstrates a fluctuation in the feedback response over the course of learning.

The major role of reflexes is to respond when there is an unexpected perturbation. In recent work Scott (2012, 2004) proposes that the majority of the sophistication embodied in the feedback stretch response occurs during the long latency component and not the short latency component. To assess whether the change in response of the triceps muscle as participants adapt to novel FF perturbations is specific to the long latency component of the stretch response, we calculated the mean triceps EMG within three time windows relative to the onset of the force pulse, -50-0 ms (PRE), 20-45 ms (SLFR), and 50-100 ms (LLFR). These measures were computed for baseline trials, early adaptation, late adaptation and washout trials. Using a two-way repeated-measures ANOVA, we compared the mean triceps EMG as a function of force pulse epoch (PRE, SLFR, and LLFR) and phase of learning (baseline, early adaptation, late adaptation, and washout). We observed a main effect of force pulse epoch (F(1,34) = 619.63, p = 2.08e-23), a main effect of phase of learning (F(1,34) = 1051.9, p = 3.68e-27), and a force pulse epoch by phase of learning interaction (F(1,34) = 638.06, p = 1.30e-23). **Figure 4** qualitatively demonstrates that there was no difference in mean triceps EMG between baseline, early adaptation, late adaptation or washout in the PRE or SLFR epochs. A *post hoc* analysis showed that during the LLFR epoch, mean EMG increased in early adaptation (2.82 ± 0.10) relative to the baseline (2.06 ± 0.04) phase of learning (t(34) = -7.10, p = 9.9299e-08, paired t-test). The effect size for the analysis (*d* = 1.2006) was found to exceed Cohen’s (1988) convention for a very large effect (*d* = 1.2). There was no reliable difference in mean triceps EMG activity during the LLFR epoch during late adaptation or washout, relative to baseline. This further demonstrates that the change in feedback response during force field learning is specific to the LLFR epoch.

To obtain a single measure of the LLFR and track changes over the course of learning, for every pair of excitatory and inhibitory force pulse trials we computed the difference between the EMG signals when we applied a 3.5 N force pulse (triceps inhibition) and EMG when a 7 N force pulse was applied (triceps excitation). We then averaged this difference signal between a 50-100 ms post-pulse window to generate a single value, termed the delta LLFR. **Figure 5** shows the time course of the delta LLFR, averaged across participants. When the force field was abruptly turned on, we observed a transient rise in the LLFR that quickly returned to baseline.

**Figure 6A** shows a summary of EMG activity during baseline trials (32 non-probe trials) and the last half of the adaptation block (56 non-probe trials) for the biceps, posterior deltoid and pectoralis. The Figure shows the across-participant mean EMG (± SEM) of each muscle, respectively, for reaching trials in the absence of force pulses or force channels. While the force pulse trials were specifically designed to inhibit or excite the triceps muscle, a response was also observed in the other three muscles recorded in this experiment (i.e., biceps brachii, pectoralis major, and posterior deltoid).

**Figure 6B** summarizes the EMG of these muscles in response to force pulses that excited (purple line) or inhibited (yellow line) the triceps lateral head. The Figure shows the across-participant average EMG (± SEM), averaged across all force pulse trials in the experiment. Because the data have all been normalized to the background load applied in the start position, the values of EMG start at 1. While the force pulse trials elicited a response in each of the other three muscles, these responses did not show the same pattern of change during adaptation as for the triceps.

**Figure 6C** shows the time course of the delta EMG for the other three muscles recorded in this experiment, averaged over the same LLFR time window as shown for triceps in Figure 5. Given that the force pulse trials were specifically designed to inhibit or excite the triceps muscle, the time course shown for the delta EMG in the biceps, pectoralis, and deltoid is only a measure of the response generated in each respective muscle during the LLFR epoch. Here the delta EMG measure represents the difference between trials in which the triceps was excited and inhibited. Qualitatively, in all three muscles, the time course is relatively flat. When we compare the average delta EMG during baseline (last four force pulse trials) and early adaptation (first four pulse trials), there is no reliable change for the biceps (p = 0.1252), pectoralis (p = 0.2298), and deltoid (p = 0.1547, paired t test). At the beginning of learning, participants experienced large errors and one possible strategy of the motor system could be to upregulate the feedback gain of all arm muscles as a mode of robust control. However, the lack of change across these three muscles demonstrates that the change in the LLFR during learning is specific to the muscle involved in countering the environmental perturbation (i.e., the triceps lateral head).

To rule out the possibility that the observed differences in triceps LLFR may be due to the muscle being stretched to a different degree in different phases of learning, we tested for differences in hand displacement during the LLFR time window (50-100 ms post force pulse). To test for differences in hand displacement during the LLFR epoch we performed a one-way repeated measures ANOVA, which indicated no reliable difference in mean hand displacement across phase of learning (p = 0.3307, **Figure 6D**). This rules out the possibility that observed changes in LLFR may be due to differences in muscle stretch.

We used a two-state model (Smith et al., 2006) to decompose the measured adaptation indices into a fast and a slow learning process. We used a procedure previously described in Coltman et al. (2019) to fit the model to data averaged across participants. The four model parameter estimates are fast retention (Af = 0.2943), fast learning rate (Bf = 0.2075), slow retention (As = 0.9992), and slow learning rate (Bs = 0.0291). **Figure 7A** shows simulated learning curves generated using the model parameter estimates. The model explains 89% of the variance in the across participant average adaptation index values over the course of learning (R^2^ = 0.89, p = 2.3143e-31). When we examine the time course of the triceps LLFR, it closely resembles the fast component of the feedforward system. To assess the similarity of the LLFR time course during the adaptation phase of learning (trials 51 - 250) and that of the fast process, we used linear regression. During the adaptation phase we have 38 data points for the delta LLFR (38 probe trials that stretched triceps and 38 probe trials that inhibited triceps). We found the corresponding value of the simulated time course for the fast process for each data point of the LLFR (i.e., value of estimated fast process at the trials at which we obtained LLFR measures) and in **Fig 7B** we show these independent probes of feedforward and feedback components together. Each data point represents one trial during adaptation, averaged over participants. Results of the Spearman correlation indicate that there was a reliable positive association between the measures of the feedforward and feedback systems: Rs^2^ = 0.88, p = 3.53 e-21. Note that while there appear to be 3 points most significantly contributing to this relationship, they should not be considered outliers, rather they correspond to the rise in the time course of both measures early in force field adaptation. **Figure 7C** shows the time course of the fast process (estimated from force channel data), overlaid on the time course of the measured triceps delta LLFR, demonstrating their remarkable similarity. We also examined the relationship between delta LLFR time course and the overall learning curve from the model output. The Spearman correlation coefficient was statistically reliable but smaller in magnitude than that for the LLFR and the fast process (Rs^2^ = -0.65, p = 7.32 e-05).

**Figure 8A** shows perpendicular deviation for all reaching (non-probe) trials, averaged across participants. To assess performance and learning during adaptation to the CCW force field, we computed the across participant mean absolute perpendicular deviation over the first four trials or last four trials during each phase of learning (**Figure 8B**). For the adaptation phase, the mean absolute perpendicular deviation in the early adaptation (first 4 trials; 1.95 ± 0.08) epoch was larger than in the late adaptation (last 4 trials; 0.57 ± 0.05) epoch (t(34)=12.66, p = 2.0040e-14, paired t-test). Characteristically the fast process allows the motor system to respond to large changes in the environment (Smith et al., 2006). To assess whether the LLFR is similarly sensitive to large errors, we regressed the values of the LLFR (the last two values in baseline and first 12 in adaptation), with the values of the perpendicular deviation on the nearest reaching trial (**Figure 8C**). This chosen window allowed us to evaluate whether the LLFR was modulated by recent errors. Each data point is the average value of each measure, at a particular trial, averaged across participants. Perpendicular deviation and delta LLFR were significantly correlated (Rs^2^ = -0.41, p = 0.0124). Again, it is important to note that while 2 data points appear to drive this relationship, they should not be considered outliers, rather they represent the first two measures taken during early force field adaptation. This suggests that LLFR is modulated by recently experienced error.

By linking the LLFR to the fast process, we predict that the LLFR would be unchanged by slower changes in the environment—changes that involve small errors, and thus would not engage the fast process. To test this idea, we performed a control experiment in which the perturbation was introduced according to a gradual adaptation schedule (**Fig. 9A**). In this way, we could train participants to reach in the same environment after the perturbation is fully ramped on, however participants would only experience small errors trial-to-trial. Qualitatively, the time course of the delta LLFR in the triceps muscle is relatively flat (**Fig. 9B**). When we compare the average response of the triceps LLFR across the epochs of learning (baseline, early adaptation, late adaptation, and washout) there is no reliable difference (p = 0.986; **Fig. 9C**). In **Fig. 9D** we compared the mean perpendicular displacement produced by the force pulses as a function of the phase of learning (baseline, early adaptation, late adaptation, and washout) and found no reliable differences (p = 0.249). Importantly, in **Figure 9E**, we show that the degree of adaptation at the end of learning was not different when the force field was introduced abruptly (Experiment 1) or gradually in the control experiment (p = 0.931). By having participants learn to adapt to the same force field as in Experiment 1, only this time using a gradual perturbation schedule, we show that this form of learning was not associated with changes in the LLFR. This finding supports previous research suggesting that learning from smaller errors engages a different process than learning from large errors that occur during abrupt perturbation schedules (Criscimagna-Hemminger et al., 2010; Schlerf et al., 2012; Tseng et al., 2007).

## Discussion

We compared changes in feedback responses to changes in feedforward control during the adaptation of reaching movements to a velocity-dependent force field. We used the time course of changes in the LLFR as a characterization of how the feedback response changes during adaptation. Feedback responses were upregulated early in adaptation and returned to baseline levels once movement errors were reduced and reaching performance reached asymptotic levels. The observed change in feedback gain was specific to the long-latency epoch and was observed only in the triceps muscle — the muscle which was required to counter the force field during adaptation. We used a two-state model of short-term motor adaptation to decompose learning into independent fast and slow processes. Overall, our findings show that feedback responses are dependent on task dynamics, they are increased in the early stages of learning, and they are reliably correlated with the fast component of feedforward adaptation.

Force field learning is a standard and often utilized experimental model of error-based motor learning (Shadmehr and Mussa-Ivaldi, 1994). The fastest changes in reaching behavior occur in the earliest stages of learning, when large errors are reduced. This initial process happens on a fast time scale, and the results of the present study show that changes in LLFR occur on a similarly rapid time scale. This may be contrasted with the literature on visuomotor rotation learning, in which a link has recently been proposed between model-based adaptation processes operating on a fast time scale and experimental probes of explicit cognitive strategies for learning (McDougle et al., 2015; Taylor et al., 2014). Visuomotor learning is driven by errors signaled by visual inputs, while errors in force field learning involve perturbations to the limb, muscle stretch and associated afferent inputs. It may be that the upregulation of feedback gains on these proprioceptive and somatosensory inputs contribute to rapid reduction of movement errors in the earliest stages of force field learning, rather than explicit cognitive strategies, which may be responsible for reducing errors on a fast time scale in visuomotor adaptation.

Additionally, we recognize that there is evidence in other forms of motor adaptation that feedback changes can occur over a slower timescale. For instance, recent work by Maeda et al. (2018) tracked the LLFR in a shoulder muscle in response to force pulses that created pure elbow motion while participants performed single-joint elbow movements, with the shoulder either free to rotate or mechanically fixed. In this case the LLFR was modulated on a much slower timescale than that observed in the present study. It’s worth noting that in the present study the experimental task was designed to probe kinematic error-based learning, while in Maeda et al. (2018) the shoulder fixation task presumably involves a very different kind of adaptation driven not by kinematic error signals but perhaps by energetic considerations.

Many design features of the present study are based on Ahmadi-Pajouh et al. (2012), but one salient difference is the use of a background load in the present study to control for the state of the muscle across the task. This enables us to attribute perturbation-related changes in EMG to learning-related changes in feedback gain, rather than changes that may be due to differences in the state of the muscle prior to movement onset. The design of the present study is considerably different from the experimental task used by Cluff and Scott (2013). In their experiment, adaptation to a velocity-dependent resistance to a single joint (the elbow) occurs at one target, while the force pulse used to probe for changes in feedback responses occurs for movements to a different target location. It is possible that this separation between the movement used for adaptation and the (different) movement used for probing feedback influenced the pattern of feedback changes observed over the course of adaptation.

The results of the present study, taken together with previous studies, demonstrate that feedback responses can be modulated according to task demands (Ahmadi-Pajouh et al., 2012; Cluff and Scott, 2013; Franklin et al., 2012; Maeda et al., 2018). In the present study changes in feedback gains over the course of learning provide insight into how the internal estimate of the dynamics of the environment is formed and used for control. By comparison, the work by Maeda et al. (2018) and Cluff and Scott (2013), both using a robotic exoskeleton, perturbing isolated joints, may provide insight into how the internal estimate of the limb, rather than the environment, is formed during learning. Studying how feedback changes differ across a range of experimental paradigms could provide valuable insights into the organization of feedforward and feedback control for motor learning.

Franklin et al. (2012) recently demonstrated that the magnitude of a rapid visuomotor feedback response is increased by both the introduction and the removal (after adaptation is complete) of a velocity-dependent force field. Furthermore, they related the size of the visuomotor feedback response to the size of perpendicular error in the reach trajectory. In the present study, by linking perpendicular error and the change in the LLFR, our results from Experiment 1 are consistent with the idea that the change in the LLFR is modulated by recently experienced error.

To test the idea that changes in the gain of the LLFR during adaptation are dependent upon experiencing large movement errors, we conducted a second experiment using a gradual perturbation schedule. Previous research suggests that learning from small, even imperceptible errors during a gradual perturbation schedule engages a different neural process than learning from large errors that occur in abrupt perturbations schedules, such as in Experiment 1 (Izawa et al., 2012; Orban de Xivry et al., 2011; Schlerf et al., 2012). We modified the schedule of the velocity-dependent curl field to test whether feedback responses change when errors are very small. Results from the control experiment showed no reliable change in the LLFR, even though participants adapted fully to the curl field by the end of the gradual schedule. This further supports the idea that changes in the gain of the LLFR are specific to large movement errors.

In recent years, researchers have increasingly asked whether feedback responses adapt when we learn new motor skills (Ahmadi-Pajouh et al., 2012; Cluff and Scott, 2013; Franklin et al., 2012; Maeda et al., 2018; Wagner and Smith, 2008; Yousif and Diedrichsen, 2012). Although these studies have shown that feedback responses can be modified with learning by increasing sensory feedback gains, how this adaptation takes place and what other features of learning feedback changes are linked to, have been largely unexplored. In the present experiment by probing feedback and feedforward components of adaptation throughout learning we were able to examine the time course of each. The time course of changes in the LLFR was strikingly similar to the time course of the fast component of feedforward adaptation. This is highly suggestive of the idea that feedforward and feedback control processes may be supported by similar learning mechanisms and neural circuits.

Orban de Xivry and colleagues (2011) have shown that primary motor cortex (M1) may play a significant role in adapting to an abrupt, but not a gradual, schedule of perturbation. Shadmehr and Krakauer (2008) propose that M1 is responsible for regulating feedback gains used to update an internal model. Pruszynski et al. (2011) have shown that M1 responds to proprioceptive feedback and that the timing of this response is consistent with the idea that it contributes to the long latency component of the feedback stretch response. Taken together, these findings suggest that the motor system can generate task-specific feedback starting as early as 50 ms following the onset of a perturbation. Feedback responses during this long-latency epoch are thought to involve cortical, brainstem, cerebellar, and spinal circuits (Cluff et al., 2015; Marsden et al., 1976; Pruszynski and Scott, 2012; Scott, 2012).

Recently Sarwary et al. (2018) probed the excitability of M1 during force field adaptation by measuring motor-evoked potentials (MEPs) in response to single-pulse trans-cranial magnetic stimulation (TMS). Their results demonstrated that the modulation of MEPs over the course of learning was correlated with the fast learning process. There is convincing evidence that the cerebellum is involved in the adaptation of predictive feedforward commands based on sensory prediction errors (Criscimagna-Hemminger et al., 2010; Schlerf et al., 2012; Smith and Shadmehr, 2005; Tseng et al., 2007). Moreover, Schlerf et al. (2012) suggest that during the rapid early phase of learning, when an abrupt perturbation is first introduced, the contribution of the cerebellum to adaptation is greatest. Taken together with the present results, these findings are consistent with the idea that during force field adaptation, the cerebellum and M1 regulate both the fast component of feedforward control, as well as the gain of the long-latency feedback stretch response.

Optimal feedback control provides a framework for us to understand how the motor system should handle performance errors caused by noise or environmental perturbation (Crevecoeur et al., 2014). Our results provide strong support for the task dependency of feedback within the framework of optimal feedback control theory. In line with previous findings, we show that there is a link between feedforward and feedback control. This supports the idea that a key feature of adaptation is to adjust feedback responses according to task demands (Ahmadi-Pajouh et al., 2012; Diedrichsen et al., 2010; Franklin et al., 2012; Wagner and Smith, 2008).

Recently Diedrichsen et al. (2010) suggested that feedforward and feedback control are not separate processes, but rather lie on a continuum. Similarly, Thoroughman and Shadmehr (2000) found that error-driven feedback produced in response to a novel force field perturbation gradually shifted over the course of learning from a feedback-driven mode of control to more predictive, feedforward control. Comparably, it has been proposed that a simple feedback control policy can be formed by modifying feedforward control with a feedback component that cancels out movement errors, thereby aiming to keep the movement as close as possible to the planned trajectory (Haith and Krakauer, 2013; Kawato and Gomi, 1992). The transient rise and fall of the LLFR time course observed in the present study suggests that as participants became faster and more accurate as a function of practice, they subsequently became less reliant on feedback control.

## Acknowledgements

Canadian Institutes of Health Research (CIHR) and the Natural Sciences and Engineering Council of Canada (NSERC). The authors thank Rodrigo Maeda for his helpful advice and comments.

*The authors declare no competing financial interests.*

